# scSPLAT, a scalable plate-based protocol for single cell WGBS library preparation

**DOI:** 10.1101/2021.10.14.464375

**Authors:** Amanda Raine, Anders Lundmark, Alva Annett, Ann-Christin Wiman, Marco Cavalli, Claes Wadelius, Claudia Bergin, Jessica Nordlund

## Abstract

DNA methylation is a central epigenetic mark that has diverse roles in gene regulation, development, and maintenance of genome integrity. 5 methyl cytosine (5mC) can be interrogated at base resolution in single cells by using bisulfite sequencing (scWGBS). Several different scWGBS strategies have been described in recent years to study DNA methylation in single cells. However, there remain limitations with respect to cost-efficiency and yield. Herein, we present a new development in the field of scWGBS library preparation; single cell Splinted Ligation Adapter Tagging (scSPLAT). scSPLAT employs a pooling strategy to facilitate sample preparation at a higher scale and throughput than previously possible. We demonstrate the accuracy and robustness of the method by generating data from 225 single K562 cells and from 309 single liver nuclei and compare scSPLAT against other scWGBS methods.

**Motivation:** scWGBS library preparation in a one-cell-per-library format presents practical and economical constraints to the number of cells that can be analyzed in a research project. In addition, most of the current scWGBS methods suffer from low read alignment rates. We present a scWGBS protocol which mitigates these issues, empowering single-cell DNA methylation analysis at an increased scale.

## Introduction

Cytosine DNA methylation is an epigenetic modification that plays a key role in the multilayered, regulatory connection between the genome and the transcriptional output in a cell. The role of DNA methylation is multifaceted and context dependent and it is known to be important for development, X-chromosome inactivation, imprinting and for genome stability. Whole Genome Bisulfite Sequencing (WGBS) methods can interrogate the methylation status of the majority of the cytosines in the genome, although DNA methylation in vertebrates occurs mostly in CpG-context (Lee et al., 2010). However, a limitation of ‘bulk’ WGBS is that DNA fragments derived from thousands of cells are randomly sampled and incorporated into the sequencing library. Consequently, with ‘bulk’ methods any observed epigenetic heterogeneity within a given sample will be difficult to interpret. Current methods for analyzing whole genome DNA methylation in single cells (scWGBS) produce sparse data. At most, the methylation status of 20-40 % of the CpG sites per cell can be measured and frequently only 5-10 % of the sites are covered (1-2 million CpG sites in human cells) (Farlik et al., 2015; Hui et al., 2018; Luo et al., 2018; Smallwood et al., 2014). Nevertheless, this level of CpG coverage is often sufficient for cell type clustering and assessment of epigenetic heterogeneity across cells (Farlik et al., 2015; Hui et al., 2018; de Souza et al., 2020). Moreover, the advent of new laboratory methods for single-cell methylation analysis have sparked an active development of computational approaches to deal with the sparsity and to impute missing data (Kapourani and Sanguinetti, 2019; Kapourani et al., 2021; de Souza et al., 2020; Tang et al., 2021). By combining scWGBS data from a handful of single cells it is possible to reconstitute near full methylomes for a cell population (Farlik et al., 2015; Hui et al., 2018) and this may prove to be especially useful when studying methylation signatures of rare cells, cells in complex tissue or clonal populations in cancer.

A variety of protocols for scWGBS library preparation have been published in the recent years. The majority of those relies on FACS sorting of single cells into 96 or 384-plates. The pioneering scWGBS method was built on the ‘post bisulfite adapter tagging’ (PBAT) method, which is used in combination with several rounds of pre-amplification of the bisulfite converted DNA prior to library preparation (Miura et al., 2012; Smallwood et al., 2014). Additional scWGBS protocols applying variations of the PBAT theme have since been published (Gravina et al., 2016; Hui et al., 2018). In a second category of scWGBS methods (Luo et al., 2017, 2018) the bisulfite converted template is subjected to a single round of strand copying and concomitant 5’-end adapter tagging. A low complexity sequence tag is then enzymatically appended to the 3-end of the DNA fragment. Although the details of the commercial Adaptase method is not disclosed, the tag functions as a handle which enables ligation of the second adapter. A third and technically distinct method for scWGBS library preparation uses a combinatorial indexing approach to barcode DNA fragments in single cells prior to bisulfite conversion. By using transposomes loaded with indexed adapter-sequences that are depleted in cytosines (thus not affected by bisulfite conversion), DNA fragments are labelled in a cell-specific manner, sequenced as a pool of several hundreds of cells, and a low number of CpG sites per cell can be interrogated after cell demultiplexing (Mulqueen et al., 2018).

Regardless of the recent progress of methods for scWGBS, compared to scRNA-seq there are few robust methods available for single cell DNA methylome analysis. The existing protocols present specific technical advantages and limitations in comparison to each other. For instance, PBAT based methods that employ several rounds of pre-amplification can interrogate up to 40 % of CpG sites in a single cell when sequenced to near saturation (Smallwood et al., 2014). However, these methods frequently suffer from chimeric reads and unacceptably low mapping efficiencies that result in high sequencing costs, which has hampered widespread adoption. The snmC-Seq2 approach typically generates data with higher mappability than other methods, but it uses commercial Adaptase reagents which results in a significantly higher cost and lower flexibility. Hence further developments in the space of single cell methylation sequencing are warranted.

We have previously developed Splinted Ligation Adapter Tagging (SPLAT), a low input method for ‘post bisulfite’ adapter tagging of single stranded DNA (Raine et al., 2017). Herein we describe single cell SPLAT (scSPLAT), as a new solution for scWGBS library preparation. The scSPLAT protocol takes cells sorted into 384-plate wells as input and after cell-barcoding and pooling of cells, SPLAT-ligation is then carried out in a bulk reaction for scalable and cost-efficient library preparation. We demonstrate the method by sequencing the methylomes of several hundreds of single cells from the K562 cell line and from a human liver sample. Our data show that scSPLAT is a robust method that compares well to existing methods and overcomes the frequent issue of low mappability of scWGBS reads.

## Results

### A SPLAT solution for single cell WGBS library preparation

Previously described scWGBS methods have used PBAT techniques or low complexity tailing of single stranded DNA 3’-ends to append Illumina adapters ‘post bisulfite conversion’ (Luo et al., 2018; Smallwood et al., 2014). We reasoned that splinted ligation could be employed in a similar manner, however with the advantage that splinted ligation applied to barcoded DNA fragments pooled from many cells could enable a more robust and cost efficient scWGBS library preparation approach. To demonstrate the scSPLAT protocol (Figure 1) we first isolated single cells or single nuclei by fluorescence-activated cell sorting (FACS) into 384-plate wells containing lysis buffer. The lysed single cells were then subjected to low-volume bisulfite conversion in 384-plate format (Luo et al., 2018). Second-strand synthesis was then performed in each cell/well using a combination of oligos (comprised of a randomer at the 3’ end, an inline cell barcode, and 26 bp of the Illumina P5 adapter at the 5’-end) and Klenow polymerase. We used random octamers of H-bases (A, C, and T) because omitting G nucleotides in the randomer was previously shown to reduce adapter dimer formation for the comparable snmC-Seq2. Exonuclease I was used to remove excess strand synthesis oligos. Thereafter, up to 32 5’-tagged second-strand reactions from the single cell reactions were pooled and SPRI-bead purified. Next, heat denaturation was performed to separate the second strand from the original strand and 3’-end adapter ligation was performed in a bulk reaction using SPLAT (Raine et al., 2017). In the SPLAT reaction, the 20 bp initial 3’ part of the P7 Illumina sequence was ligated to the 3’-end of the previously tagged and barcoded single strand DNA molecules, using splinted dsDNA adapters with a protruding random hexamer the 3’-end of the bottom strand, which acts as a splint and hybridizes with the single stranded DNA, enabling ligation using T4 DNA ligase (Figure 1). Finally, the pooled and twice adapter tagged fragments were PCR amplified with unique dual index primers containing the remainder of the Illumina barcode sequences for 12-16 cycles, which generated > 2 nM of sequencing library, which was sufficient for quality control, pooling and sequencing on an Illumina NovaSeq sequencer (see Figure S1 for examples of library profiles). After sequencing, cell-demultiplexing was performed based on the inline cell barcode which is the first six bases of read 1.

**Figure 1.**
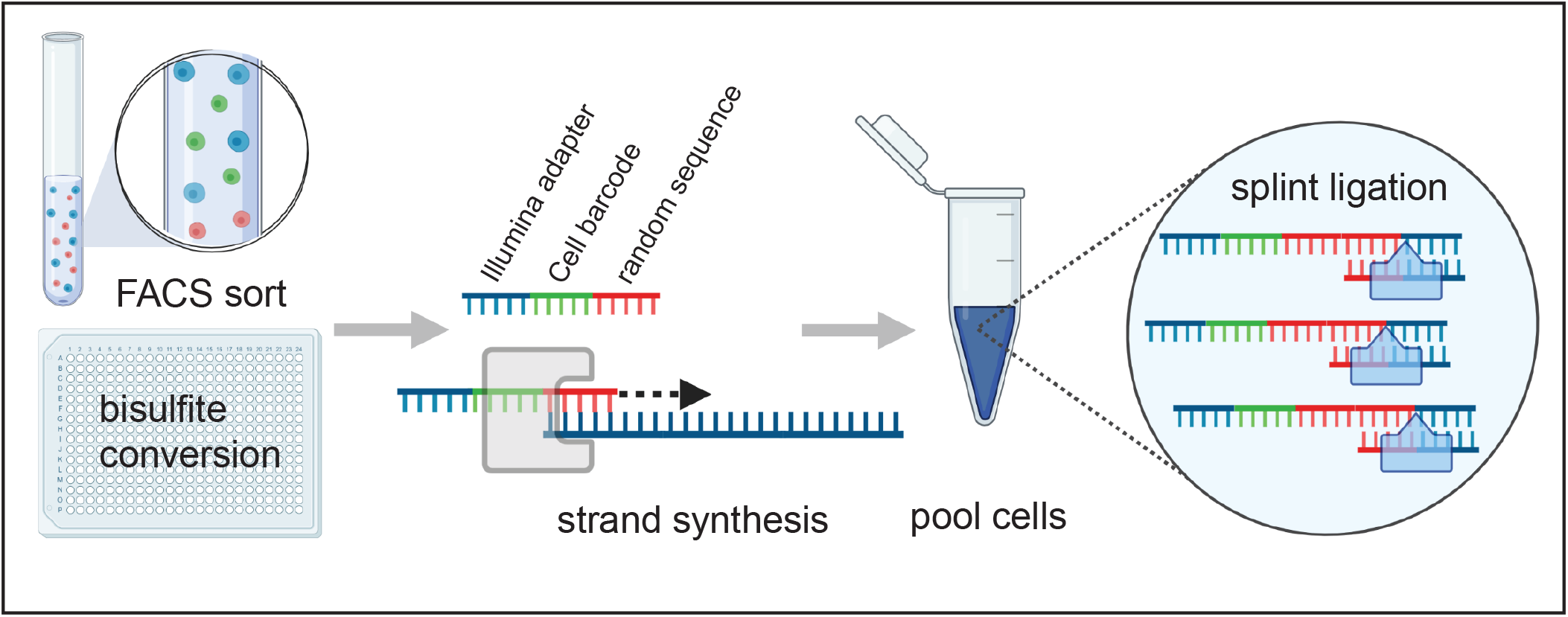
Schematic overview of the scSPLAT method. A) Cells are FACS sorted into 384 wells containing lysis buffer and bisulfite conversion is performed in 384-plate format. B) A random strand synthesis reaction is performed in each well using a randomer flagged with an inline cell barcode and 26 bp of the Illumina P5 adapter sequence. C) Reactions from multiple wells (up to 32) are then pooled and SPRI bead purified. D) Splinted ligation adapter tagging (SPLAT) is performed to ligate the adapter to the 3’-ends of the DNA fragments in a bulk reaction.

### Single cell SPLAT applied to the leukemia K562 cell line

To evaluate scSPLAT, we first prepared and sequenced 225 single K562 cells prepared in two separate batches. The scSPLAT libraries were prepared either as 8- or 16-single-cell pools. (Figure 2A). In addition, we prepared 16 single K562 cells (2×8-cell pools) using a comparable method (snmC-Seq2). We sequenced 2-60 M read pairs (PE150) per cell for scSPLAT libraries and 7.5-20 M read pairs/cell for the snmC-Seq2 libraries. More than 99% of reads in the pools could be assigned to cells based on the inline barcode. We observed no apparent overrepresentation of specific cell barcodes in the pools and the fraction of reads assigned per cell were relatively even (Figure S2).

**Figure 2.**
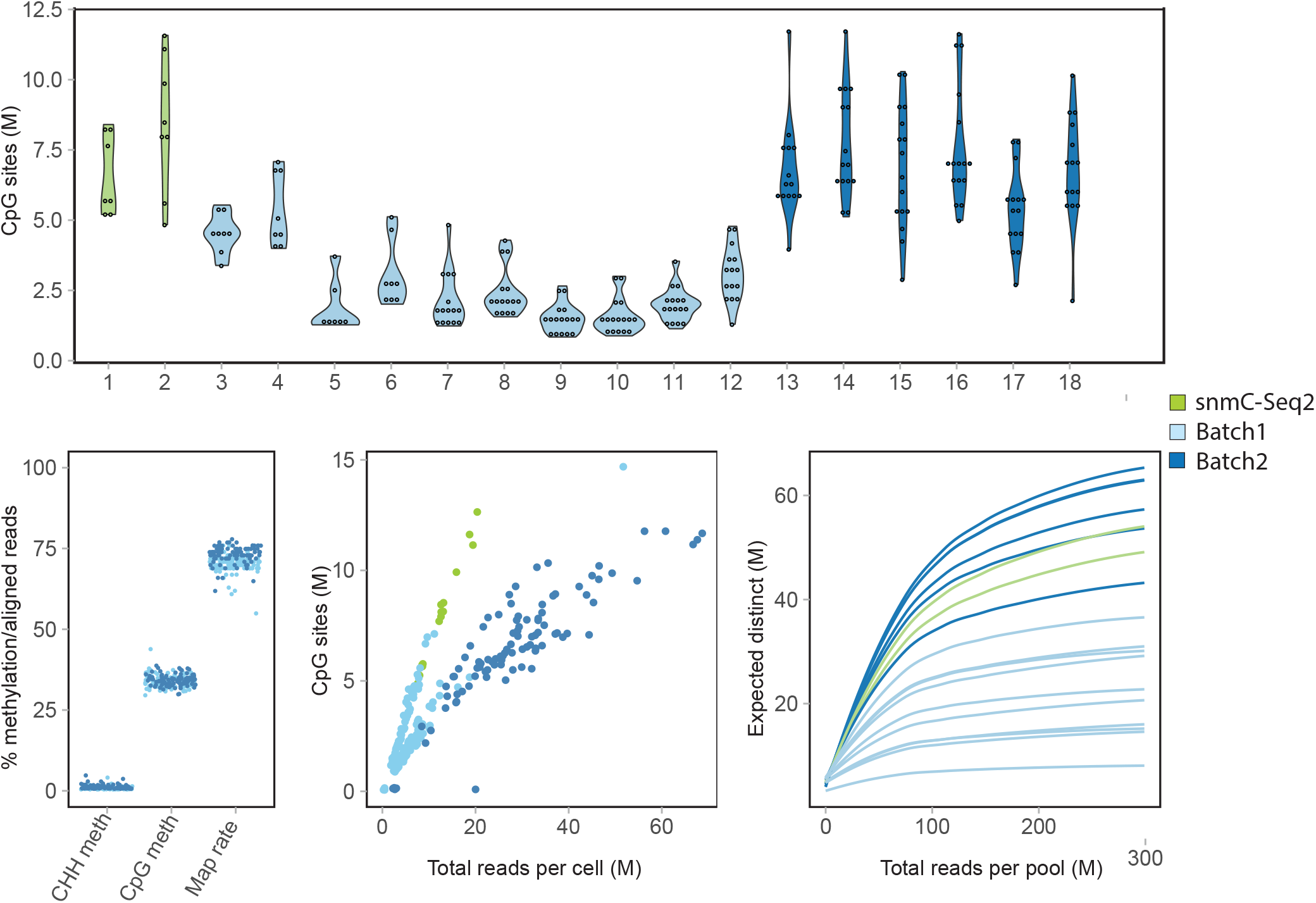
Quality assessment of single cell K562 data. A) Violin plots showing the total number of covered CpG sites/cell in each library pool. B) The percentage of CHH methylation (indicative of bisulfite conversion efficiency), methylation levels in CpG-context and read mapping efficiency per K562 cell. C) The number of CpG sites per cell covered ≥ 1x plotted as a function of the total number of (raw) reads generated per cell. D) Pool-wise library complexities plotted with the c-curve function in the preseq tool.

Poor mappability is a common issue with PBAT derived scWGBS reads and frequently only as low as 20-40 % of the reads align to the reference (Gravina et al., 2016; Hui et al., 2018; Smallwood et al., 2014). In contrast, other methods like e.g snmC-Seq2 tend to perform better (> 50%) with respect to mapping efficiency. This is presumably due to the single round of strand copying, which likely reduces the likelihood for non-mappable chimeric and artificial sequences to be formed. Similar as for snmC-Seq2, only one strand synthesis reaction is performed in the scSPLAT protocol and therefore high mappability of reads was expected. We applied a read mapping procedure wherein we: i) mapped the reads paired-end (pe), ii) re-mapped the unmapped fraction as single-reads (sr), and iii) combined the pe and sr mapped data. Indeed, using this strategy the mean percentage of reads mapping unambiguously to the GRCh38 reference was 71.6 +/-3.01 for scSPLAT, thus approaching numbers normally observed for bulk WGBS (Figure 2B). Similarly, when using the same approach for the snmC-seq-2 data, the mean mapping efficiency was comparable at 70.5 +/-1.89 (Data S1).

In total, 94 % of the scSPLAT K562 cells passed stringent quality filtering, in which we removed cells with lower than 50 % mapping rate, higher than 2% CHH methylation rate (indicative of inefficient bisulfite conversion), and cells with < 500,000 CpG sites covered (by 1 sequencing read). The average methylation rates per cell in CpG and CHH context are shown in figure 2B. The resulting 211 single cell methylomes were then used for the downstream analysis and clustering.

The number of strand-merged CpG sites covered by ≥ 1 read ranged between 1-11.5 Million/cell for the K562 cells prepared with scSPLAT, i.e up to 40% of all CpG sites in the human genome (Figure 2A). In the snmC-seq2 control libraries 5-12 Million CpG sites/cell were covered. As expected, the CpG site coverage was higher in cells with higher sequencing depth (Figure 2C). However, we also observed substantial variance in library complexity across pools, where library pools from the first batch of libraries generally reached saturation at lower sequencing depth compared to the second batch of libraries, as well as the snmC-Seq2 data (Figure 2D).

To validate the methylation levels obtained with scSPLAT data against previously generated methylation data for the same cell type, we downloaded two bulk K562 WGBS data sets from ENCODE (average CpG-site coverage 30x and 35x) (Dunham et al., 2012; Zhang et al., 2020). We then merged the quality-passed scSPLAT cell-libraries batch-wise into data sets with average CpG coverage of 20x and 30x (pe+sr mapped) for batch 1 and batch 2, respectively and computed pairwise correlations across all sets (including CpG sites 5 ≥ x coverage). The Pearson’s correlation was consistently high when comparing to ENCODE data (R =0.93-0.94) as well as for comparisons across the batches (R=0.97) (Figure 3A) indicating accurate methylation levels measured by scSPLAT.

**Figure 3.**
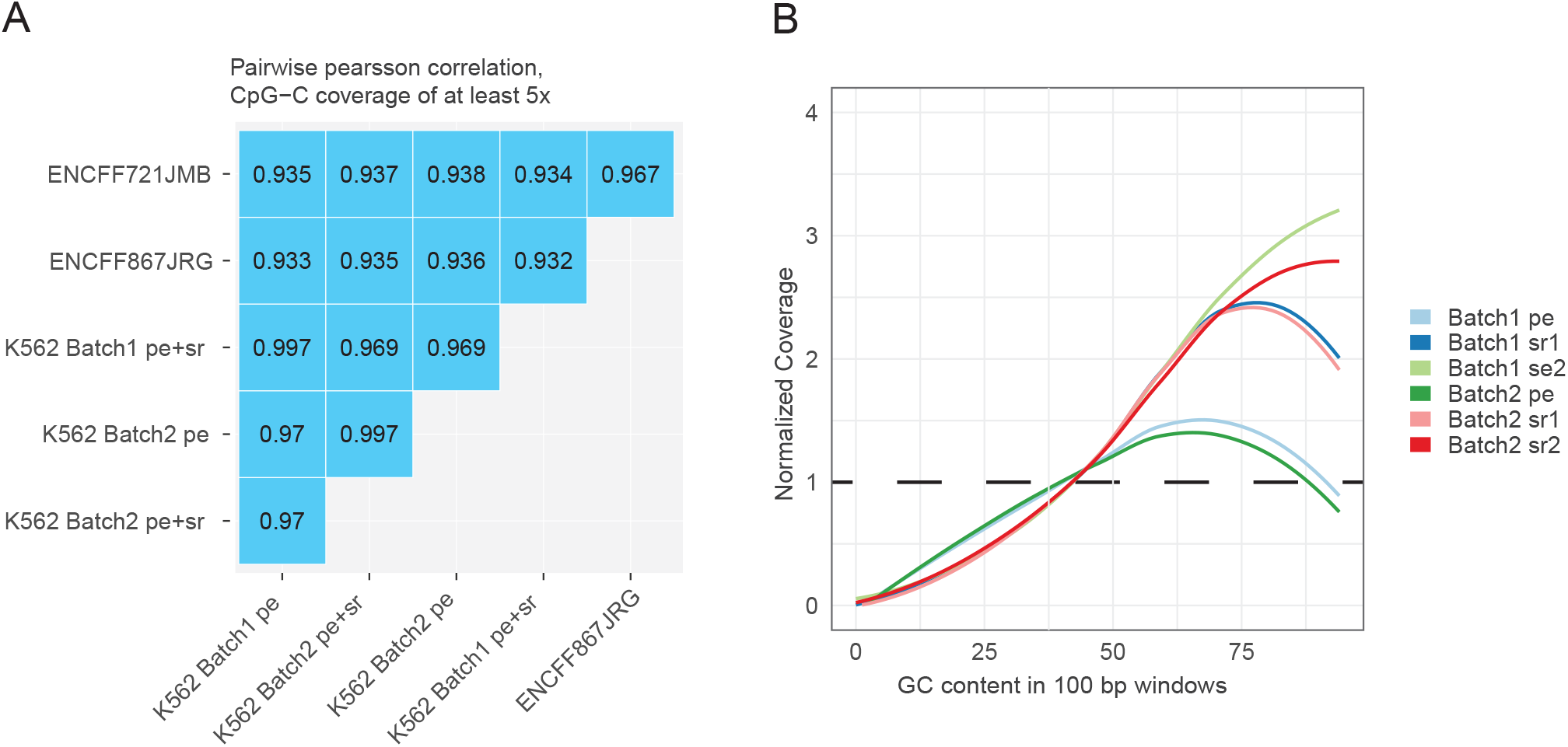
Assessment of methylation levels and GC bias in pseudo-bulk scSPLAT K562 data. A) Single cell data were merged batch-wise to produce ‘pseudo’ bulk data sets. Pairwise methylation correlation was computed across the pseudo-bulk SPLAT data and two sets of K562 ENCODE bulk WGBS data. Pearson’s R values are shown for comparisons across ENCODE sets and pseudo-bulk SPLAT mapped in paired-end (pe) mode as well as in combined paired end and single read mode (pe+sr). B) GC bias profiles for batch-wise merged pe and sr mapped reads respectively. Coverage was shifted towards genomic regions of higher GC content, especially for the sr mapped data (reads that were unmappable in pe mode and subsequently mapped as single reads).

Next, we also investigated coverage bias by analyzing read coverage in regions of varying GC content for the two batch-wise merged scSPLAT K562 data sets. Similar to what has been shown for previous scWGBS methods (Farlik et al., 2015), coverage appears to be positively biased toward genomic regions with higher-than-average GC content. Notably, the GC-bias was more pronounced for the single end mapped reads suggesting that a higher fraction of those reads map to GC rich repeat regions (Figure 3B).

To explore underlying structures in the scSPLAT data we employed the Epiclomal framework (de Souza et al., 2020) for probabilistic clustering of cells according to their methylation profiles. As an additional control we downloaded a set of K562 scWGBS data generated in a previous study (Farlik et al., 2015). Epiclomal Region applied across CpG-islands followed by UMAP-visualization grouped K562 cells prepared with different methods (scSPLAT cells, snmC-Seq2 cells and Farlik-scWGBS) into a single cluster as would be expected for a homogeneous cell-line sample (Figure 4A). Moreover, cells derived from the same library pool did not cluster together, although there was a tendency for cells from the two scSPLAT batches to separate within the cluster (Figure S3). This seeming batch effect is likely explained by a generally lower CpG site coverage in CpG islands (due to both shallower sequencing and lower library complexity) in batch 1 compared to batch 2 (Figure 2D, data S1). We also performed clustering based on CpG sites in transcription start site (TSS) regions and gene bodies (GB) resulting in the same single cluster (Figure S4). The visual batch effects in the TSS and GB based clustering appeared subtler compared to the CpG island clustering (Figure S4). Taken together, these data demonstrate that the scSPLAT method correctly and robustly assesses DNA methylation in single K562 cells.

**Figure 4.**
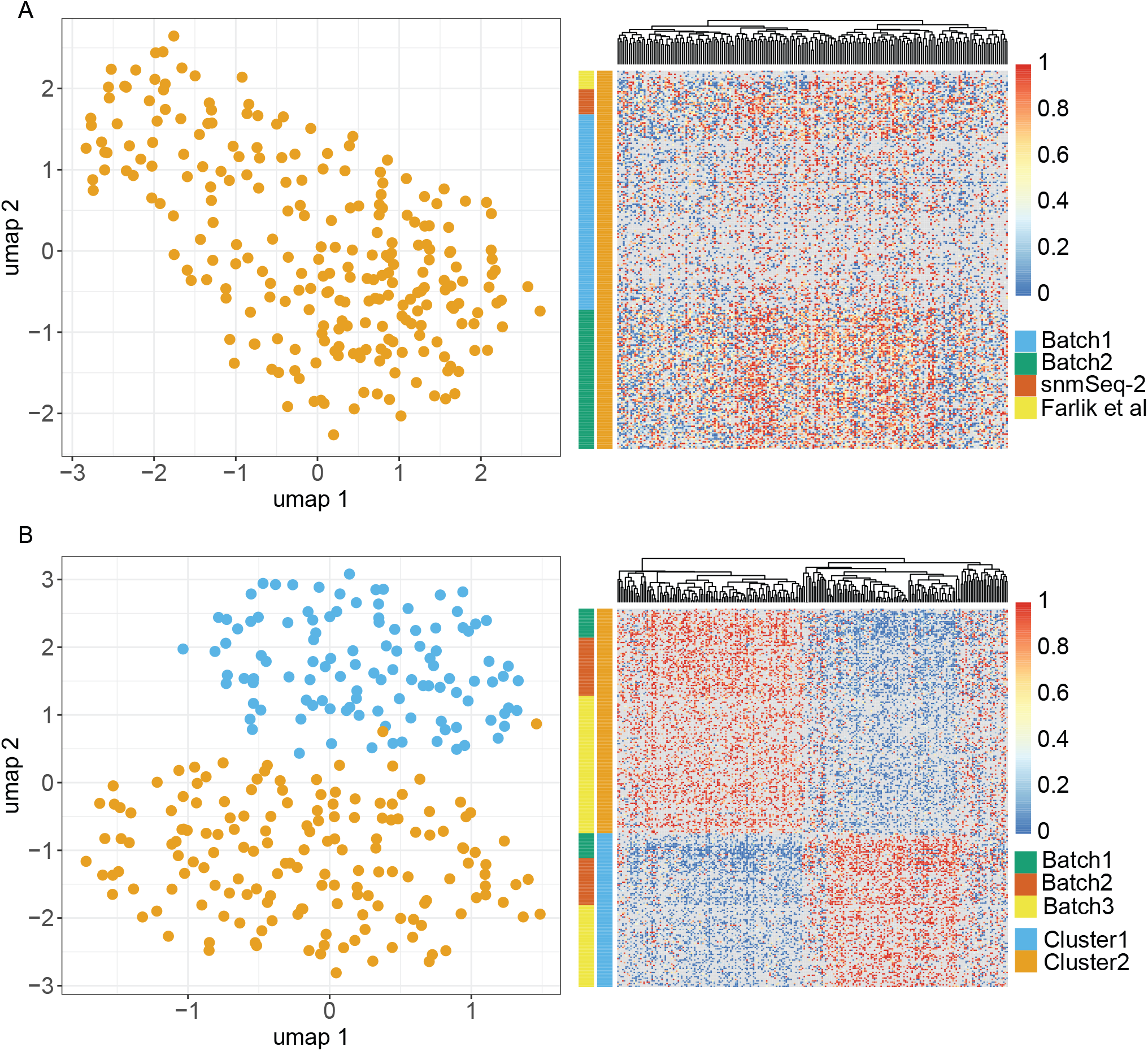
Clustering of K562 and Liver nuclei based on single cell DNA methylation profiles. A) UMAP visualization of Epiclomal Region clustering of K562 cells based on CpG-island methylation. B) Heat map showing mean methylation of the regions (CpG-islands) used as input for Epiclomal Region clustering of K562 cells. Blue color indicates no methylation and red color high methylation, Greys boxes in the heatmap indicates missing values. C) UMAP visualization of Epiclomal Region clustering of liver nuclei based on CpG-island methylation. D) Heat map showing mean methylation of the regions (CpG-islands) used as input for Epiclomal Region clustering of liver nuclei.

### scSPLAT applied to 309 single nuclei from a human liver sample identifies two major clusters

To validate scSPLAT in a more complex sample type and to evaluate scSPLAT on nuclei, we next applied the method to 309 single nuclei isolated from a human liver sample (Diamanti et al., 2021). The library pools comprised either 4, 16 or 32 cells from the same specimen and were prepared on three different occasions (batches 1-3). Similar as in the previous experiments with K562 cells, demultiplexed reads were evenly distributed across cells in the pools (Figure S2). Using the same computational approach for mapping the reads as for the previous K562 experiments, a mean value of 68 +/-5.62 % of liver nuclei reads mapped unambiguously to the human reference genome (Data S1). After quality filtering, 276 nuclei remained for downstream clustering analysis and 0.6-4.7 million CpG sites/cell were detected by at least one read. The global CpG methylation rate for the liver nuclei ranged between 66-82% (Figure S5).

The human liver represents a heterogeneous, but well-defined tissue where the cell population is dominated by hepatocytes. Other major cell types present in the liver are non-parenchymal cells such as e.g hepatic stellate cells and liver sinusoidal endothelial cells (Kmiec, 2001). Previous studies investigating transcriptomes of single nuclei obtained from the same specific liver sample used herein identified hepatocytes followed by hepatic stellate cells and endothelial cells as the most common cell populations in the sample (Cavalli et al., 2020; Diamanti et al., 2021). When applying the Epiclomal Region clustering method to the 273 liver nuclei, we observed two major clusters/cell populations (Figure 4B). Nearly identical cluster assignments were obtained irrespective of whether Epiclomal Region was applied with methylation values across CpG-islands, gene bodies or TSS regions (Data S2). Importantly, nuclei derived from different batches and pools were evenly distributed throughout the clusters implying little batch effect (Figure S3).

For certain cell types, e.g neurons, there is a relatively well-defined association between gene activity and CH-context methylation in gene bodies (Liu et al., 2020; Uzun et al., 2021). However, for most other cell-types, gene body methylation in CH-context is very low and the relationship between CpG-methylation and gene activity is typically less clear, posing challenges to classification of cell-types based on marker-gene associated DNA methylation. Nevertheless, we attempted to cell-type label the two clusters. First, we pooled the methylation information from nuclei cluster-wise and then performed differential methylation analysis across the two merged data sets. Differentially methylated regions (DMRs) were defined as comprising at least 10 CpG sites and having an absolute difference in mean methylation > 0.6 between cluster 1 and 2. The DMRs were annotated to genomic regions and to the nearest gene (Data S3). This resulted in a total of 5114 DMRs. The majority of DMRs were in introns or intergenic regions, 349 were located in annotated promoters and 1,246 were found within +/-2000 bp of a transcription start site (TSS). The number of annotated ‘nearest genes’ (hereafter denoted DMR-genes) were 3779, and 901 of the DMR-genes were annotated to two or more DMRs. A subset of the DMR-genes with cluster specific hypomethylated DMRs in or near promoters are shown in Figure 5.

**Figure 5.**
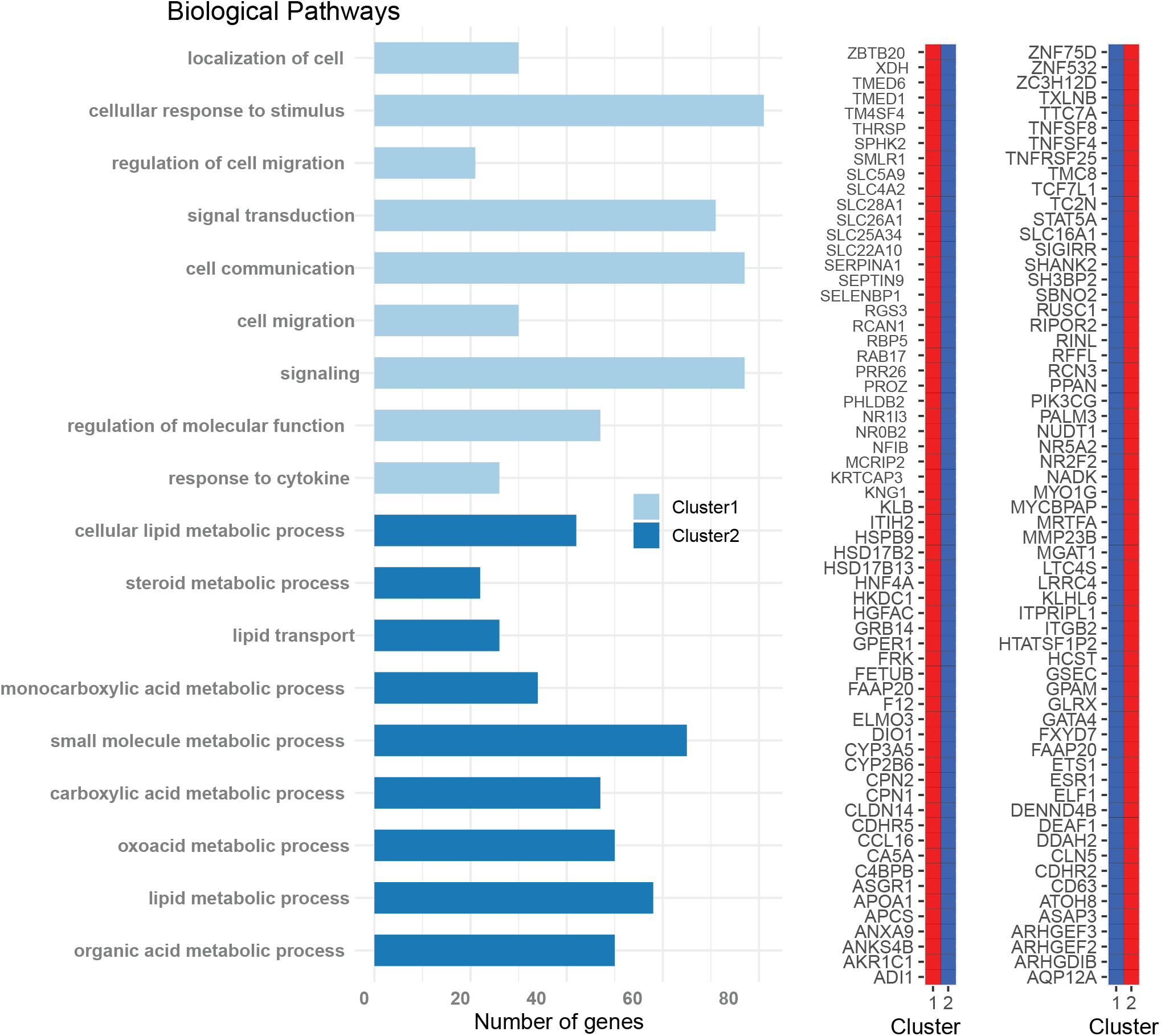
Cluster specific gene ontology pathway analysis. A) Genes associated with a hypomethylated DMR in cluster 1 or cluster 2 and located in a promoter and/or TSS region, were used as input for GO pathway analysis. The most significant biological pathways found for cluster 1 and cluster 2, respectively are shown. Metabolic pathways were identified in cluster 2, indicating hepatocytes (HCs). For cluster 1 the identified GO pathways did not reveal a specific cell type B) A subset of the genes with DMRs in promoter regions. The DNA methylation status of the DMRs in each respective cluster are shown (low methylation = blue, high methylation = red)

If a DMR was hypomethylated in one of the clusters and was in a promoter region, CpG island, or within 500 bp upstream or downstream of a TSS site, its DMR-gene was included in a test for Gene Ontology (GO) biological pathway enrichment-This resulted in 182 and 385 DMR-genes for GO analysis of cluster 1 and cluster 2, respectively. The GO pathway analysis of cluster 2 resulted in a clear enrichment of hepatocyte-specific metabolic pathways for the genes with hypomethylated DMRs in the larger cluster, revealing that the identity of the 163 nuclei in cluster 2 is hepatocytes (Figure 5 and Data S4). To corroborate accurate classification of cluster 2, we next sought to investigate if genes known to be hepatocyte cell-markers were associated with the DMRs identified in this study. First, we selected genes with cell-specific expression in hepatocytes as determined by scRNA-seq analysis by the Human Protein Atlas (HPA) consortium (n=190 genes). (Karlsson et al., 2021). Second, we selected the genes used as hepatocyte markers from a previous study snRNA-seq study that included the same liver sample analyzed herein (n=76 genes) (Cavalli et al., 2020; Diamanti et al., 2021). Next, we interrogated the overlap between established hepatocyte-marker genes and the DMR-genes identified by scSPLAT. We found that all of hepatocyte marker-genes identified by both HPA and Cavalli *et al* had at least one, sometimes several cluster-2 hypomethylated DMRs in their vicinity, supporting our annotation of cluster 2 as hepatocytes (Data S5).

GO pathway analysis of the 182 cluster-1 hypomethylated DMR-genes displayed a less distinct profile, showing significant enrichment of pathways involved in signaling, cell communication and response to stimuli (Figure 5, Data S4). Previous snRNA-seq studies in the same liver sample analyzed herein identified hepatic stellate cells and endothelial cells next the two most abundant cell types after hepatocytes (Cavalli et al., 2020; Diamanti et al., 2021). It was therefore our hypothesis that cluster 1 (n=111 cells) is comprised of hepatic stellate cells and/or endothelial cells. We attempted to clarify the identity of the cluster by repeating the aforementioned annotation procedure with hepatic stellate- and endothelial-marker genes. In this analysis, we used the genes listed as cell-specific and group-enriched by HPA (n=144 stellate and n = 76 endothelial cell markers) (Karlsson et al., 2021). Twenty stellate cell markers overlapped with a DMR-gene, although the DMRs were not consistently hypomethylated in cluster 1 (n=11 hypo- and n= 9 hyper-methylated). Notably, four of the overlapping, hypomethylated stellate cell markers (*DNASE1L3, EHD3, TBXA2R, LIFR*,) were among the thirty genes that were classified as cell-specific by HPA i.e having > 4-fold higher expression level than any other cell type. Their corresponding DMRs were located in intergenic regions or introns. Only eleven endothelial-markers overlap with a cluster 1 DMR (n= 6 hypo and n= 5 hyper-methylated) none of those were endothelial cell-specific and thus only weakly suggest an endothelial cell contribution to Cluster 1.

The cluster 1 DMR-genes overlapped with nineteen hepatic stellate markers (wherein only three DMRs were hypomethylated) and eleven endothelial markers (n= 6 hypo and n= 5 hyper-methylated) identified in the study by Cavalli *et al* (data S5). Thus, the DMR-overlap with markers from that study did not further clarify the cell-type composition of cluster 1.

To investigate if the resolution of clustering could be improved, we applied Epiclomal Region to the 111 cells in cluster 1. However, this did not result in any substructure that could clearly indicate whether there are one or two sub-populations in cluster 1. (Figure S6). In summary, we observed a number of DMRs associated with genes shown to specifically have increased expression in hepatic stellate cells (Karlsson et al., 2021). However, it was not possible to conclusively label cluster 1 with a specific cell-type based on the DNA methylation profiles. The results herein demonstrate that cell types with very specialized functions, such as hepatocytes, are easier to annotate based on their DNA methylation profiles, while other cell types (such as hepatic stellate and endothelial cells) are more difficult to annotate on DNA methylation profiles alone.

## Discussion

In recent years there has been a surge of experimental protocols for gene expression analysis and epigenetic features in single cells. scWGBS is a robust alternative for assessment of cell-to-cell epigenetic heterogeneity and may also be useful for discovery of previously unknown cell types/states. One advantage is that DNA methylation is less influenced by common confounders known to pose problems to single-cell RNA-seq analysis, such as stochastic expression, transcriptional bursts, and cell cycle effects. However, it is technically challenging to analyze DNA methylation at the single cell level, hence large-scale single cell methylome studies are still in their infancy. Nevertheless, one striking example of the utility of scWGBS is in a recent study that profiled the methylomes of > 100,000 mouse brain nuclei using the snmC-Seq2 method (Liu et al., 2021). To facilitate single cell methylome analysis on a scale larger than a few hundred cells, easily implemented library preparation methods in combination with pooling strategies will be essential. Moreover, as sequencing costs continue to drop, inexpensive library preparation methods become more attractive. The scSPLAT approach presented herein is a robust and cost-efficient protocol which confers higher read map rates compared to most other scWGBS methods and allows for pooling of cells for higher throughput without the need of using a commercial kit. Furthermore, as an open protocol it is flexible, can be tailored for project needs and further optimized for improved performance. We herein performed pooling of up to 32 cells in a single library. This number is by no means the upper limit for pooling, and larger pools will enable even lower library costs per cell. By scaling down the second strand reaction volume e.g using micro dispensing systems, we envision that full 384-plates reactions can be pooled in a single library.

scSPLAT is in several respects similar to the previously described scnmC-seq2 method, which is why we included in-house prepared scnmC-seq2 libraries as controls in the present study. In contrast to the Adaptase based scnmC-seq2 protocol, no artificial low complexity sequence is appended during adapter ligation in the scSPLAT workflow. The appendage of such low-complexity sequence tags both take up extra sequencing space and need to be carefully trimmed so as to not interfere with read mapping and methylation calling. In addition, for scnmC-Seq2 library preparation caution must be taken so that there is no carryover of free-nucleotides into the pool prior to the Adaptase reaction. It was previously shown that free nucleotides from the strand synthesis reaction could interfere with the low complexity tagging by producing artificial tag sequences of unexpected compositions. This may lead to inefficient tag trimming and in turn lower map rates and biased methylation calls. In contrast, scSPLAT adapter tagging is performed using splint ligation and carryover of free-nucleotides poses no risk for introduction of artificial sequences.

From a technical point of view the scSPLAT method performed well when applied to both to a cell line and nuclei from primary tissue. Using our *ad hoc* DMR analysis approach we found distinct signatures in cluster 2 matching hepatocytes. The cluster 1 cell population possibly comprises both hepatic stellate and endothelial cells, however our clustering approach did not separate the cell types and the DMR analysis did not yield a strong signature for either of them. Interestingly, a previous study analyzing scRNA-seq expression in various cell types showed that hepatic stellate and endothelial cells types had related RNA expression profiles (Karlsson et al., 2021) and it is therefore likely that they have similar DNA methylation landscapes. Our results corroborate the fact that it can be challenging to annotate certain cell types based on DNA methylation alone. It is possible that the non-parenchymal cells were too few and that increasing the number of cells and/or alternative clustering approaches would increase the cluster resolution. Joint gene expression and methylation profiling in the same single cell will facilitate cell type annotation (Uzun et al., 2021) and our protocol presented here can be further developed to enable this.

## Limitations of the study

Although scSPLAT confers robust measurement of DNA methylation levels (batch effects were low), there were quite some variances in library preparation efficiency across batches, observed as differences in complexity and duplication rates. There may be multiple causes for differences in library complexity; suboptimal cell lysis, low bisulfite conversion yields, strand synthesis efficiency and ligation efficiency. Further work will be required to pin-point and address the main causes of this variability. Meanwhile it might be worthwhile to perform library pool amplifications using real time PCR to avoid overamplification and to estimate library complexity based on initial low coverage sequencing and exclude low quality pools prior to deeper sequencing.

## Acknowledgements

This research received funding from the European Union’s Horizon 2020 research and innovation program under grant agreement No. 824110EASI-Genomics. The project was also partially funded by the Swedish Research Council (2019-01976), the Göran Gustafsson Foundation, and the SciLifeLab Uppsala Technology Development Grant.Sequencing was performed at the SNP&SEQ Technology Platform, part of the National Genomics Infrastructure hosted by SciLifelab and funded by the Swedish Research Council. Computational resources were provided by the Swedish National Infrastructure for Computing (SNIC) also partially funded by the Swedish Research Council. We thank the Microbial Single Cell Genomics facility at SciLifeLab for FACS sorting of the K652 cells and the BioVis facility at Uppsala University for assistance with sorting of liver nuclei.

## Author contribution

AR: conceived the method, performed lab work, analyzed and interpreted data, wrote the manuscript; AL and AA analyzed and interpreted data. ACW performed lab work and method optimization. CB performed lab work and contributed with expertise in FACS sorting. MC and CW provided FACS sorted single liver nuclei and advised on the liver nuclei cell type annotation. JN provided funding, interpreted data, reviewed and edited the manuscript. All authors have read and approved the manuscript.

## Supplementary Information

Figures S1-S6

Data S1 Sample and read alignment information

Data S2 Liver cluster assignments

Data S3 DMR annotations

Data S4 GO:BP intersections

Data S5 DMR-marker overlaps

Table S1 Oligo sequences

## Methods

### Cell sorting

Live K562 cells were sorted into 384-well plates using a MoFlo™ Astrios EQ (Beckman Coulter) with 1x PBS as sheath fluid at a pressure of 25 psi and a 100 µm nozzle. Cells were stained with Hoechst 33342 and propidium iodide (PI) (both ThermoFisher Scientific) and sorted based on forward scatter, positive fluorescence for Hoechst 33342, and negative fluorescence for PI in stringent single sort mode with a drop envelope of 0.5. Liver nuclei were sorted into 384-well plates using BD FACSMelody™ cell sorter with BD FACSChorus™ software. Nuclei were stained with 4′,6-diamidino-2-phenylindole (DAPI) and sorted using the “Single Cell” sort mode at a pressure of 23 psi using a 100 µm nozzle.

### scWGBS library preparation and sequencing

Cells were FACS sorted into 384-plate wells containing 2 µl lysis buffer (Zymo M-digestion buffer also containing proteinase K). 15 µl EZ Methylation Direct conversion reagent (Zymo Research) was added, and incubation was performed in a thermocycler according to the manufacturer’s instruction. The bisulfite conversion reaction was subsequently washed and desulfonated using reduced volumes in a 384-plate spin column (Zymo Research) essentially as described in (Luo et al., 2018) with minor modifications. Briefly; 60 µl binding buffer was first added to each well in the column plate. Then 20 µl binding buffer was added to each reaction in the 384-well plate using a multi-pipette, the reactions (37 µl) was transferred to the column plate and mixing was performed by pipetting. The spin plate was placed on top of a 96-, 2 ml deep well plate and centrifuged (5000 g, 5 min). 100 µl M-wash buffer was added and the centrifugation was repeated. 50 µl M-desulphonation buffer was then added to each well and incubated for 15 min at RT, whereafter the plate was centrifuged (5000 g, 5 min). Two washes were subsequently performed with 100 ul M-wash buffer (5000 g, 5 min). After the second wash the spin plate was placed on an empty collection plate and briefly spun to remove residual wash buffer. The spin plate was then placed on a new 384-well plate and 7 ul elution buffer containing 500 nM of the barcoded strand synthesis oligos (SSO_H_X, Table S1) was added and incubated at RT for 5 min, followed by centrifugation (5000 g, 5 min) to elute the DNA. After a brief denaturation step at 95 °C (2 min) and cooling on ice, a strand synthesis reaction was performed by adding 5 µl of Klenow mastermix (2x Blue buffer, 1 mM dNTP, 2.5 units Klenow exo-(Enzymatics) to each well and incubating at 4°C for 5 min followed by 25° for 5 min and 37°C for 60 min. Then 0.5 µl of Exonuclease I was added to each well to digest excess oligos followed by incubation at 37 °C for 30 min. After oligo digestion, up to 32 single cell reactions were pooled and 1.2x SPRI bead purified (AMPureX beads, Beckman Coulter). The DNA was eluted in 10 µl of 10 mM Tris-Cl (pH8.5) and denatured at 95°C for 2 min in the presence of 50 ng of thermostable single stranded binding protein, ET-SSB (New England Biolabs) and then immediately cooled down on ice. SPLAT was performed in bulk by adding the SPLAT 3’ adapter at 5 µM final conc, 1 µl 10x T4 DNA ligase buffer, 1 µl PEG4000 (10x stock), 15 units T4 DNA ligase (ThermoFisher Scientific) 10 units of T4 polynucleotide kinase (ThermoFisher Scientific)) and H_2_0 to a final volume of 15 µl. The ligation mix was incubated at 20 °C for 1 hr. A 1.0 x SPRI bead purification was then performed and the DNA was eluted in 10 µl. Libraries were amplified by adding 2.5 µl NEBNext dual index primers and 12.5 ul KAPA HiFi 2x PCR mastermix (Roche). PCR was carried out for 12-16 cycles and thereafter the PCR reactions were purified twice with 0.8x SPRI beads and finally eluted in 10 µl elution buffer (10 mM Tris pH 8.0). The scSPLAT pools were quantified using Qubit or TapeStation and were then combined prior to final concentration determination using qPCR. The libraries were PE150 sequenced on either HiSeqX (Liver nuclei batch 1) or a NovaSeq 6000 (Illumina). scnmC-Seq2 libraries were prepared according to the previously published protocol (Luo et al., 2018) and sequenced in the same NovaSeq lane as the scSPLAT K562 batch1 libraries. The SPLAT adapter was prepared from oligo ss1a and ss2b (Table S1) by combining 100 µM of each oligo in 50 ul, adding 1µl 20x TE buffer and 1µl of 5M NaCl and heating to 95 °C, followed by stepwise decrease to 4 °C.

### Cell demultiplexing and read mapping

Pooled scSPLAT libraries were demultiplexed into single cell samples based on the six bases long inline barcode using the je software suite v.1.2 (Girardot et al., 2016) with default arguments and specifying the barcode-presence on read 1. Prior to read alignment adapter sequences were trimmed using TrimGalore v.0.6.1 and an additional 15, resp 10 bases were clipped from the 5’- and 3’-prime end of reads for both scSPLAT and snmC-Seq2. Bismark (v.0.22.1 specifying parameters; –comprehensive, minimum alignment score function of L,0, -0.2 and maximum insert size = 800 bp), were then used to paired end align scWGBS reads to the human reference GRCh38 (Krueger and Andrews, 2011). Afterwards the unmapped fraction was re-mapped as single reads (SR) and the bam files from the respective PE and SR1 and SR2 mappings were first deduplicated and then methylation calling was performed using Bismark’s methylation extractor tool. The CpG methylation information from PE and SR mappings were combined and strand-merged using a custom R-script. Bisulfite conversion efficiency was assessed by global methylation levels in CHH context. Cells with a CHH methylation value > 2% were excluded from the analysis.

For complexity analysis the K562 pools were paired end aligned using Bismark before cell-demultiplexing. The plots were generated using the c-curve function from the Preseq software (Daley and Smith, 2013).

### Comparison of scSPLAT K562 methylation against ENCODE data and GC bias analysis

We downloaded the call sets from the ENCODE portal (Dunham et al., 2012) with the following identifiers ENCFF721JMB, ENCFF867JRG. scSPLAT bamfiles were merged using Picard MergeSamFiles and methylation levels extracted with Bismark. Common sets of CpG sites were extracted using custom R-scripts and Pairwise Pearson’s correlation coefficient were computed in R and including only CpG-site covered by at least 5x.

### GC bias analysis

The paired end and single read mapped bam files for each cell were merged into ‘pseudo-bulk’ bam files as to obtain one paired-end-, one single-read 1- and one single read 2 bam file per K562 batch. GC biases were computed using the Picard CollectGCBiasMetrics tool and normalized coverage was plotted as a function of the GC content in the genome.

### Clustering and visualization

The cells were clustered using the EpiclomalRegion tool as described by de Souza *et al*. The preprocessing pipeline provided by Epiclomal was used to construct an input matrix from the strand merged bismark .cov files and a file specifying the regions of interest. The regions selected herein were CpG islands, gene bodies and 1000 bp upstream and downstream of the transcription start sites. The region coordinates were downloaded from UCSC using custom scripts and the coordinates for the CpG sites within each region were extracted using the R package BSgenome.Hsapiens.UCSC.hg38 (v1.4.3). The preprocessing pipeline starts by obtaining the methylation levels for each CpG site within the regions from the coverage files. Non-redundant regions were identified by first filtering the regions with less than 5% coverage in 10% of cells and then inferring the interquartile range (IQR) and keeping the most variable regions. The matrix produced by the preprocessing pipeline was used as input for the EpiclomalRegion. The clustering pipeline starts by clustering the cells using four modes of non-probabilistic clustering. The results from this step are then used as a starting point together with a set of random values for the probabilistic clustering. The posterior probabilities outputted from Epiclomal were used to assign each cell a cluster. Heatmaps were plotted using the mean methylation levels across the selected regions and UMAP was used for dimensionality reduction and visualization of the cells.

### Differential methylation and GO pathway analysis

Methylation information in strand-merged Bismark.cov-files were merged cluster-wise. The R-package bsseq (Hansen et al., 2012) was used to smooth methylation calls, segment data and call DMRs between clusters. To call a segment as a DMR a minimum of ten consecutive CpG sites and an average difference > 0.6 in the DNA methylation levels between cluster 1 and 2 was required yielding 5114 DMRs. Differentially methylated regions (DMRs) were annotated to genomic regions and to the nearest gene using Homer v 4.1 (Heinz et al., 2010). The nearest genes to DMRs within promoters, CpG Islands and 500 bp upstreams and downstreams of TSS were used as input for Gene Ontology biological pathway enrichment analysis using the gprofiler2 R package (Peterson et al., 2020). Cell type specific marker genes were obtained from (Cavalli et al., 2020; Diamanti et al., 2021) and from the Human Protein Atlas; https://www.proteinatlas.org/humanproteome/celltype (Karlsson et al., 2021).

---

**Supplementary Figure 1.**
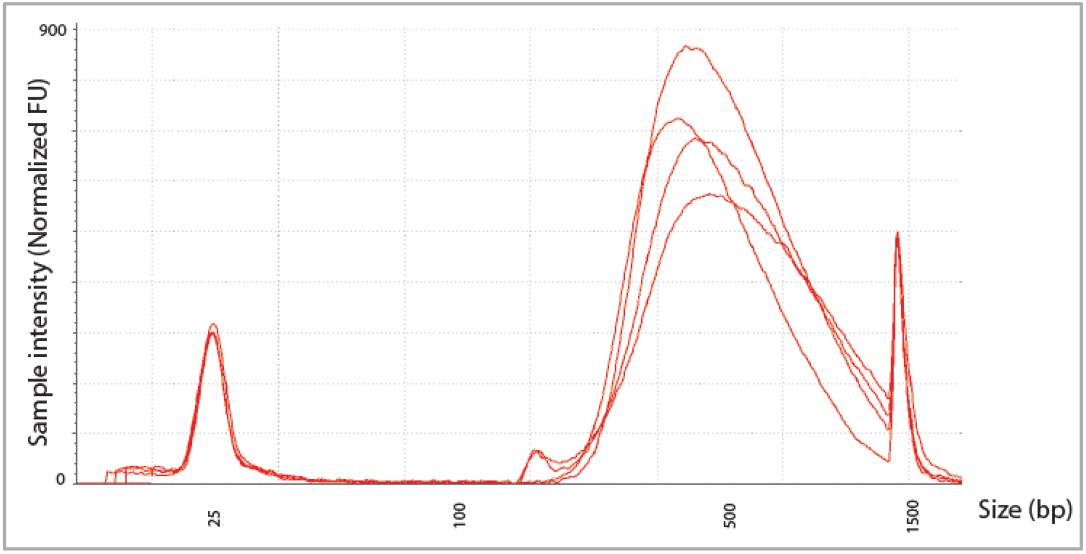
Examples of TapeStation profiles for pooled scSPLAT libraries

**Supplementary Figure 2.**
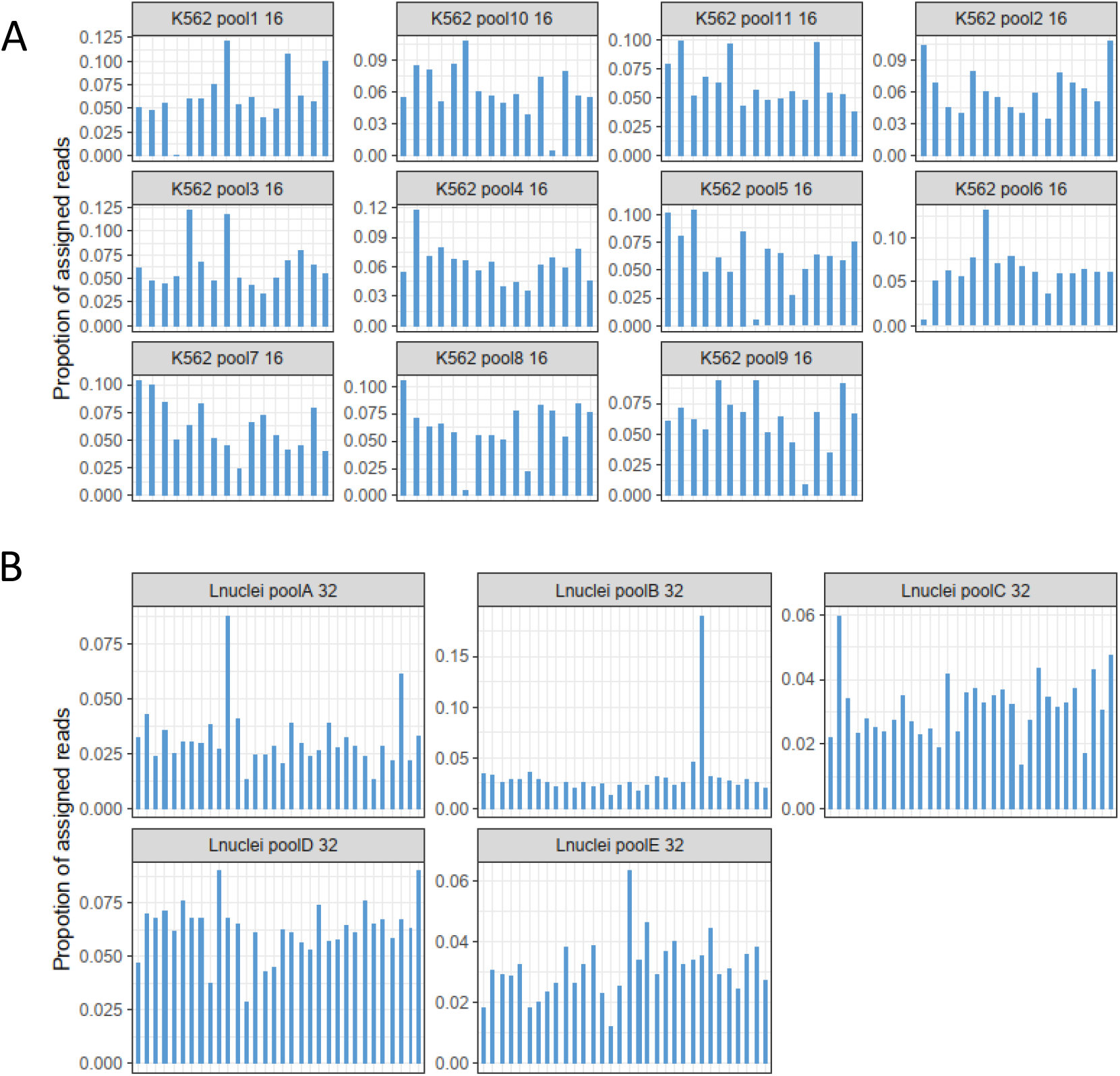
Demultiplexing profiles for A) K562 16-cell pools and B) liver nuclei 32-cell pools. The bars represent the proportion of reads assigned to each cell in the pool.

**Supplementary Figure 3.**
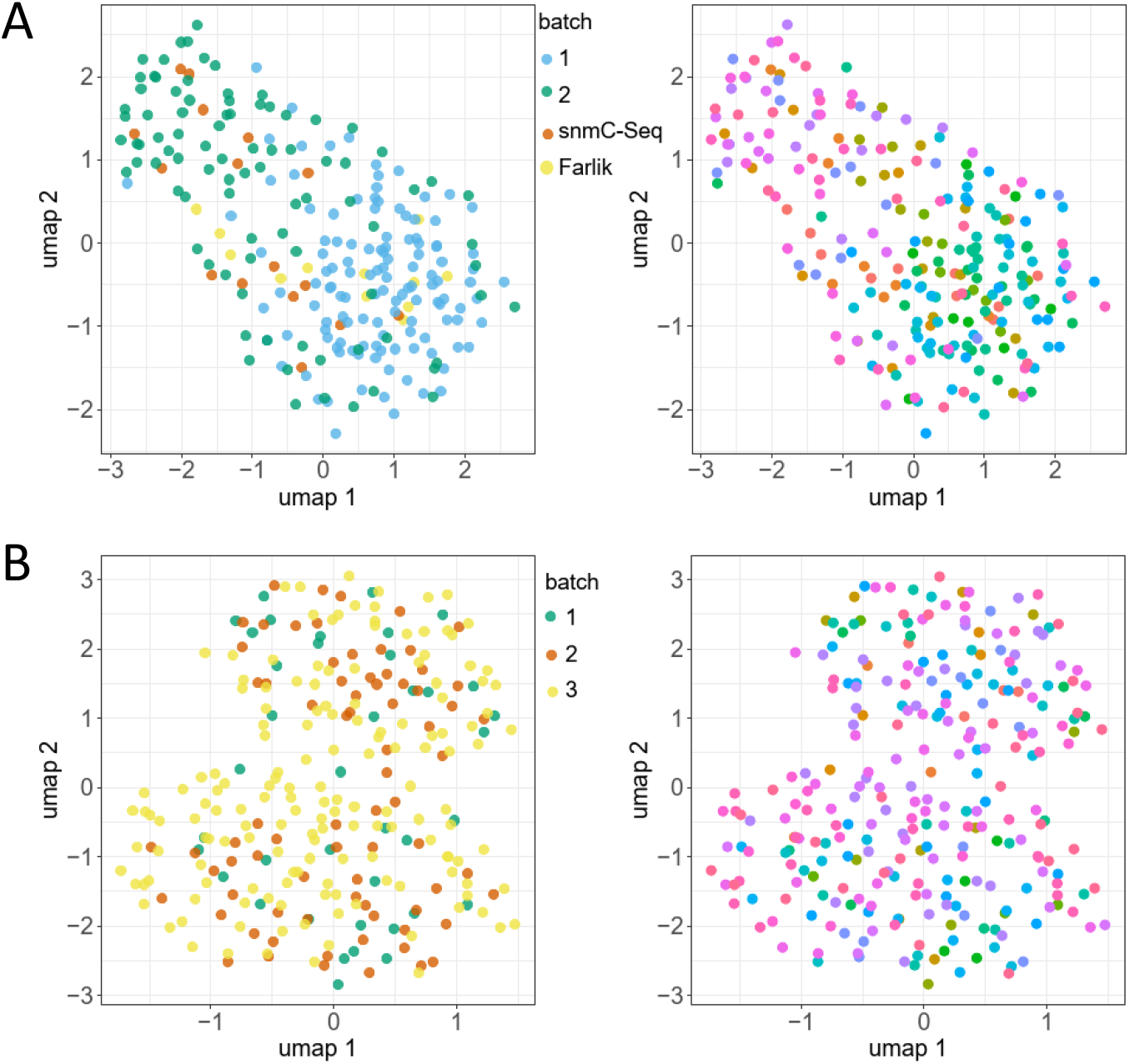
K562 and liver nuclei EpiClomal Region clustering based on methylation in CpG islands. A) K562 cells colored by batch. B) K562 cells colored pool-wise. D) Liver nuclei colored by batch. B) Liver nuclei colored pool-wise.

**Supplementary Figure 4.**
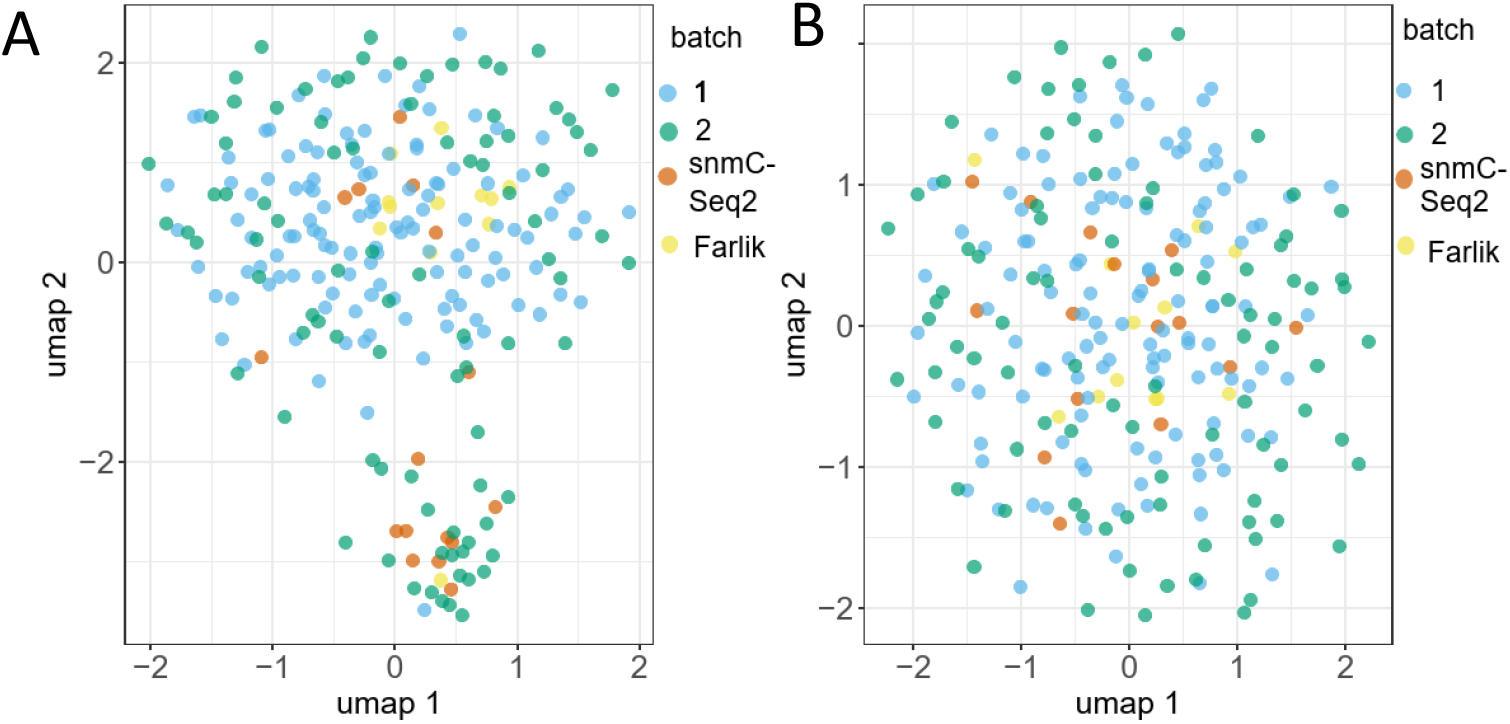
EpiClomal Region assigned all K562 cells to a single cluster irrespective of whether the clustering was performed based on methylation in CpG islands, gene bodies or TSS regions. A) K562 EpiClomal Region clustering based on methylation in gene bodies. Cells colored by batch. Although it appears like a small number of cells form a ‘subcluster’ in the UMAP plot, they were all assigned to one single cluster by EpiClomal. B) K562 EpiClomal Region clustering based on methylation in regions of 1000 bp +/- of TSS. Cells colored by batch.

**Supplementary Figure 5.**
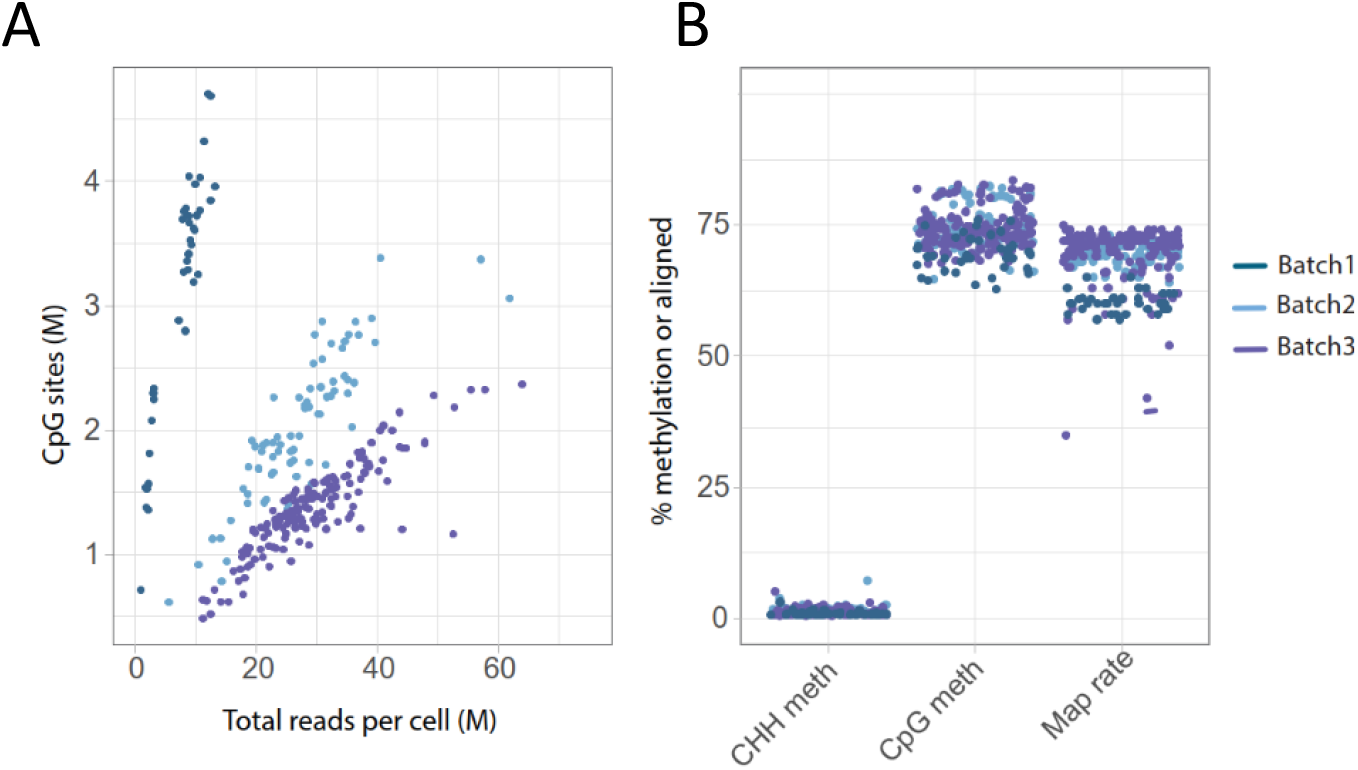
Liver nuclei QC metrics A) Number of detected CpG sites per cell vs sequencing depth B) Global methylation levels and read alignment efficiency per nuclei.

**Supplementary Figure 6.**
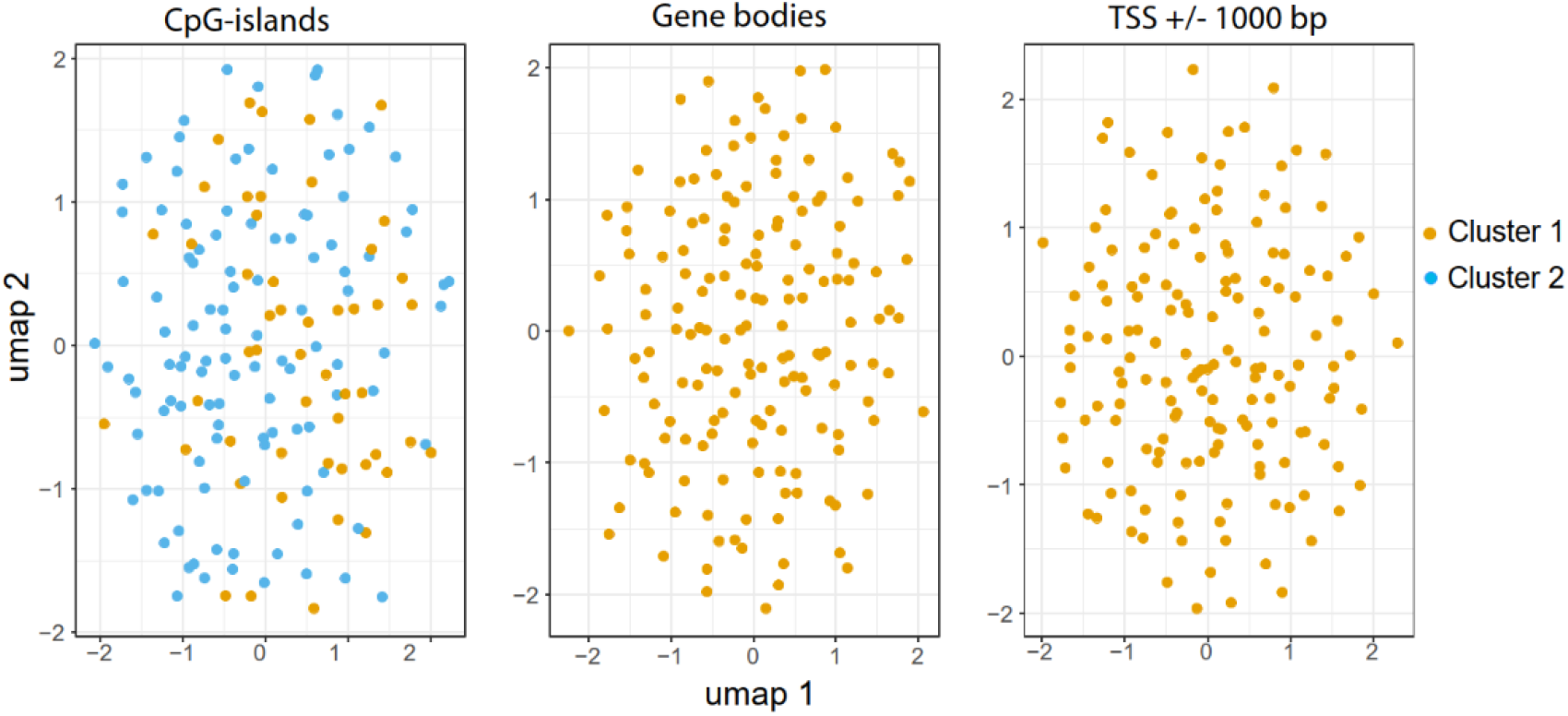
Reclustering of liver nuclei cluster 1 based on: CpG island methylation, gene body methylation and methylation across TSS regions. With the two latter EpiClomal finds one cluster only. In contrast, with CpG-island methylation EpiClomal assigns the cells to two clusters that were were intermingled in the UMAP visualization.

